# Melanopsin retinal ganglion cell-driven contribution to visual and cognitive brain responses in LHON

**DOI:** 10.1101/2020.09.04.282830

**Authors:** Stefania Evangelisti, Chiara La Morgia, Claudia Testa, David N Manners, Leonardo Brizi, Claudio Bianchini, Michele Carbonelli, Piero Barboni, Alfredo A. Sadun, Caterina Tonon, Valerio Carelli, Gilles Vandewalle, Raffaele Lodi

**Affiliations:** Unità di RM Funzionale, Dipartimento di Scienze Biomediche e Neuromotorie, Università di Bologna, Bologna, Italy; Dipartimento di Scienze Biomediche e Neuromotorie, Università di Bologna, Bologna, Italy; IRCCS Istituto delle Scienze Neurologiche di Bologna, UOC Clinica Neurologica, Bologna, Italy; Dipartimento di Fisica ed Astronomia, Università di Bologna, Bologna, Italy; Centro Fermi - Museo Storico della Fisica e Centro Studi e Ricerche «Enrico Fermi», Roma, Italy; Studio Oculistico d’Azeglio, Bologna, Italy; Doheny Eye Institute, Department of Ophthalmology, David Geffen School of Medicine at University of California, Los Angeles, Los Angeles, California, United States; IRCCS Istituto delle Scienze Neurologiche di Bologna, Programma Neuroimmagini Funzionali e Molecolari, Bologna, Italy; Sleep and chronobiology lab; GIGA-Cyclotron Research Centre/In vivo imaging, University of Liège, Belgium

## Abstract

Melanopsin retinal ganglion cells (mRGCs) are intrinsically photosensitive photoreceptors contributing to visual and non-image-forming functions of the eye. Isolating mRGC roles in humans is challenging, therefore mRGCs functions remains to be fully characterized.

We explored mRGCs contribution to light-driven visual and cognitive brain responses in Leber’s Hereditary Optic Neuropathy (LHON), given mRGC relative sparing in LHON. Twelve patients and twelve matched healthy controls (HC) participated in an fMRI protocol including visual and visual-cognitive paradigms under blue (480nm) and red light (620nm).

Higher occipital activation was found in response to sustained blue *vs.* red stimulation in LHON *vs.* HC. Similarly, brain responses to the executive task were larger under blue *vs.* red light in LHON over lateral prefrontal cortex.

These findings are in line with LHON mRGCs relative sparing and support mRGCs contribution to non-visual and visual functions in humans, with potential implication for visual rehabilitation in optic neuropathy patients.

## INTRODUCTION

Human rods and cones represent the main retinal photoreceptors of the image-forming system. However, another retinal photoreceptor system exists, relying heavily on melanopsin, a photopigment expressed in about 1% of retinal ganglion cells (RGCs) (Provencio et al., 2002). Melanopsin is maximally sensitive to blue light (~480nm) and melanopsin RGCs (mRGCs) are intrinsically photosensitive and characterized by sustained and sluggish responses to light (Berson et al., 2002; Dacey et al., 2005; Hankins et al., 2008). mRGCs are essential for the non-image-forming (NIF) functions of light, i.e. those functions of light that are not directly related to image-formation, such as circadian rhythm photoentrainment, pupillary light reflex, melatonin suppression, as well as the regulation of alertness, sleep and cognition (Gooley et al., 2012; Sand et al., 2012; Gaggioni et al., 2014). Recent evidences also support an involvement of mRGCs in visual processes, such as brightness detection and coarse image formation (Hankins et al., 2008; Ecker et al., 2010; Gooley et al., 2012; Sand et al., 2012; Gaggioni et al., 2014; Allen et al., 2019). mRGCs main central projections include the hypothalamic suprachiasmatic nucleus (SCN), site of the master circadian clock, the hypothalamic preoptic area implicated in sleep initiation, the olivary pretectal nucleus regulating pupil size, the medial amygdala, part of the olfactory and emotional response (Hattar et al., 2006; Hannibal et al., 2014). mRGCs also project to regions typically part of the visual pathway, such as the dorsal division of thalamus LGN and the midbrain superior colliculus (Hannibal et al., 2014).

Light stimulates cognitive brain activity (Vandewalle et al., 2009; Mitolo et al., 2018) and functional MRI (fMRI) studies showed that, in normally sighted individuals, light, geared towards mRGCs increases brain activity over the frontal eye field and inferior frontal cortex (Hung et al., 2017) and potentially in a region encompassing the suprachiasmatic nucleus (McGlashan et al., 2018).

Likewise, the NIF system was shown to modulate attentional, executive and emotional functions, likely through mRGCs (Chellappa et al., 2014) with maximal efficiency with blue light around 460-480nm (Gaggioni et al., 2014). However, rod and cone photoreception, contribute to mRGC light responses (Güler et al., 2008; Gaggioni et al., 2014; Chellappa et al., 2014), making the isolation of mRGC specific roles challenging in humans.

Outer retina degeneration in totally blind patients has been used as a successful model to demonstrate mRGC contribution to NIF functions (Czeisler et al., 1995; Zaidi et al., 2007; Gooley et al., 2012; Hull et al., 2018) and to evaluate the NIF impact of light on cognition (Vandewalle et al., 2013, 2018). Study samples were however small due to the rarity of the phenotype, making a generalization of mRGC signalling impact on cognition uncertain.

Leber’s hereditary optic neuropathy (LHON, estimated prevalence: 1:45,000) (Mascialino et al., 2012) is a maternally inherited blinding disorder due to mitochondrial dysfunction (Carelli et al., 2004). This is usually due to one of three mitochondrial DNA (mtDNA) point mutations (m.11778G>A/MT-ND4, m.14484T>C/MT-ND6, m.3460G>A/MT-ND1) that affect genes encoding complex I subunits (ND) of the respiratory chain (Carelli et al., 2004). In LHON patients, optic nerve atrophy occurs consequent to degeneration of RGCs in the inner retina, whereas outer retina rods and cones are preserved. Structural MR showed microstructural alterations along the visual pathway (Rizzo et al., 2012; Manners et al., 2015) and grey matter loss in the visual cortex (Barcella et al., 2010). Yet, despite the loss of regular RGCs, mRGCs are remarkably well preserved in LHON, as demonstrated by retinal *post-mortem* histopathology and *in-vivo* preservation of light-induced suppression of nocturnal melatonin secretion (La Morgia et al., 2010) and PLR (Kawasaki et al., 2010; Moura et al., 2013). Since LHON is relatively common compared to outer retinal degeneration, it provides a unique opportunity to further characterize functions of relatively preserved mRGC in the context of the severe optic nerve atrophy with increased statistical power. We reasoned that the important RGC neurodegeneration, would blunt image-forming photoreception generated by intact rods and cones, whereas preserved mRGCs NIF-image-forming and image-forming contributions would be emphasized.

We recorded brain activity of LHON patients and healthy controls (HC) in fMRI while exposed to blue (480 nm) and red (620 nm) light of different durations, in conjunction or not with a cognitive task. We could therefore explore both NIF and image-forming impacts, in the setting of disease-induced loss of regular RGCs. As for the image-forming effects, we anticipated that blue and red lights would have a similar impact on occipital pole activity in HC, while in LHON patients, in which mRGCs can represent up to 30% of the remaining RGC (as opposed to ~1% in HC), blue light would trigger larger occipital cortex activation than red light, particular for longer duration exposure (50s) compared to the short stimulation. As for the NIF impact of light in visuo-cognitive context, we anticipated that the differential impact of blue vs. red light on ongoing cognitive activity would be larger in LHON patients, with relatively more mRGCs, as compared in HC. Since MR scanner static magnetic field was lower than previous studies in HC (1.5T vs. 3T), we anticipated that difference between light conditions would be most prominent in patients.

## RESULTS

### Demographic, clinical and behavioural results

LHON patients and HC did not significantly differ in terms of age (mean ± sd, LHON: 38.2 ± 12.9 years, HC: 37.8 ± 13.7 years, t-test p=0.95), gender (M/F, LHON: 10/2, HC: 8/4, Pearson’s χ^2^ Test p=0.35) and average number of hours of light at the time of fMRI acquisitions (mean ± sd, LHON: 12.6 ± 2.1, HC: 12.7 ± 2.2, t-test p=0.95).

None of the participants had an extreme morning-evening chronotype (mean ± sd, LHON: 59.8 ± 9.3, HC: 57.4 ± 5.0, p=0.44), nor presented excessive sleep-wake disturbances, as evaluated by PSQI (LHON: 4.9 ± 2.3, HC: 3.7 ± 2.2, p=0.22), ESS (LHON: 6.8 ± 3.8, HC: 7.3 ± 1.4, p=0.65) and Berlin questionnaires. Beck anxiety and depression scores were normal in all participants (respectively LHON: 12.6 ± 7.7, HC: 5.4 ± 6.8, p=0.05 and LHON: 7.2 ± 6.5, HC: 5.9 ± 2.9, p=0.55), except for two patients who presented mild to moderate levels of anxiety and depression. Ophthalmological data of LHON patients are reported in Table 1. Fundus examination revealed a diffuse optic atrophy for all LHON participants, and Ishihara’s Test score was 0/12 for all of them. Visual field examination was not available for two LHON patients with a very severe visual loss for which the exam was not reliable. For the same reason, for 4 out of 12 LHON patients the VF of only one eye was considered for subsequent analyses. The duration of the disease in LHON patients was 17 ± 12 years and the average visual acuity for the entire cohort of LHON patients was 20/630. Average RNFL thickness, as evaluated by OCT, was 45.3 ± 5.1 microns. HC subjects had normal ophthalmological exam including OCT and visual acuity was 20/20 in all of them.

**Table 1.**
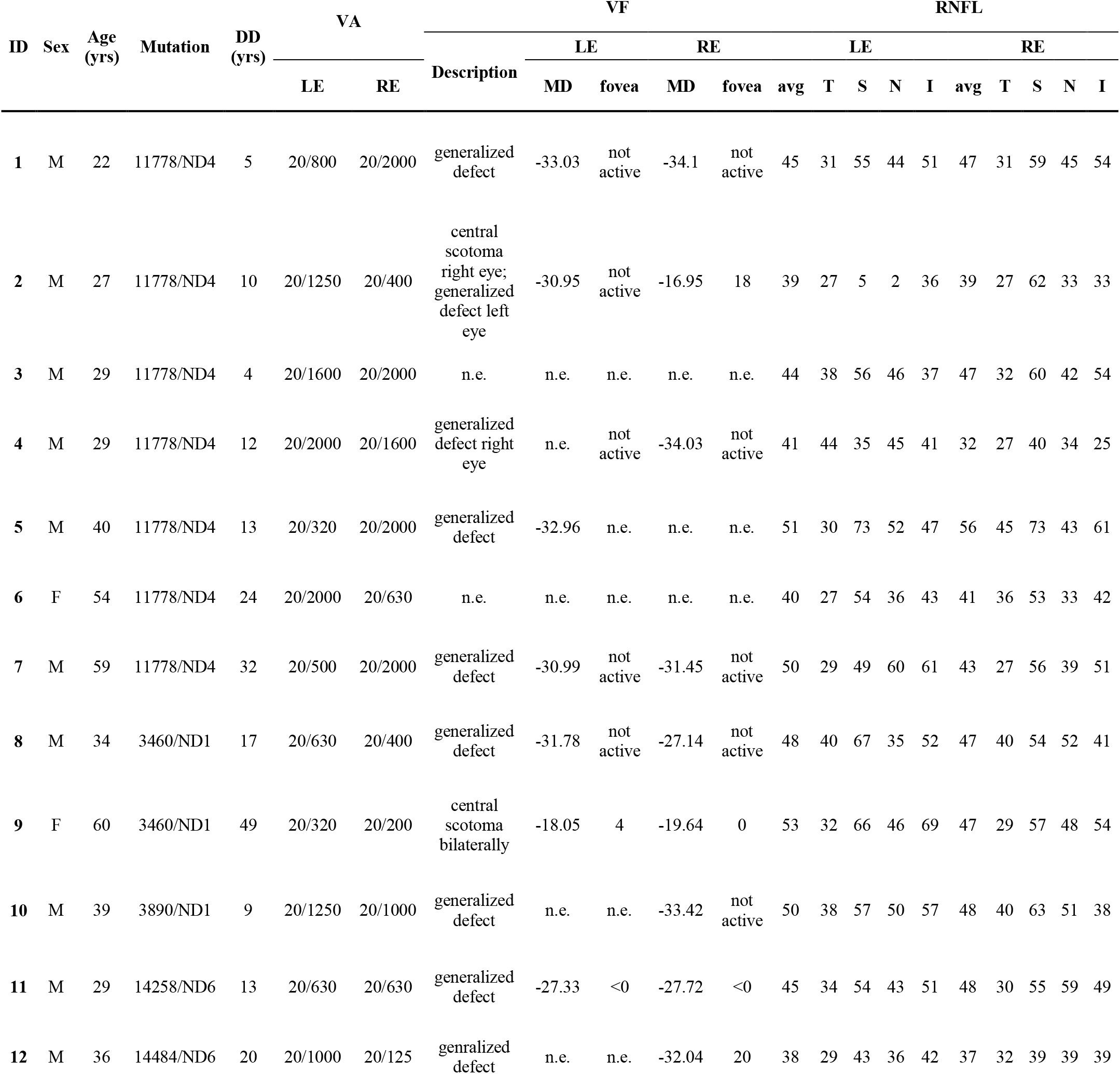
Clinical and ophthalmological evaluations for LHON patients. (DD: disease duration; LE: left eye; RE: right eye; VA: visual acuity; HM: hand motion; VF: visual field; MD: mean defect; RNFL: retinal nerve fibre layer; avg=average, T: temporal; S: superior; N: nasal; I: inferior; n.e.: not executed, due to unfeasibility).

At the second training session, all the participants reached at least 75% of accuracy in both n-back tasks. Over the whole study cohort, there was a modest but significant improvement of performances between the first training (during the week before MRI acquisitions) and the second one (just before MRI acquisitions) in the 3-back task (paired t-test, p=0.020, mean first training: 85.7%, mean second training: 89.3%).

As for the accuracy to n-back tasks during fMRI acquisition, as intended given the short block duration of both task and light exposures, there was no significant main effect of group (0-back: F=0.552 p=473; 3-back: F=0.759 p=0.402), nor light condition (0-back: F=2.861 p=0.113; 3-back: F=3.732 p=0.056), as well as no significant interaction between group and light condition (0-back: F=1.379 p=0.272; 3-back: F=1.932 p=0.174). These results imply that the fMRI results were not biased by significant differences in the cognitive task performances.

### Narrowband light stimulations

We first considered the impact of light exposure only, i.e. independent of the presence of a cognitive task. Brain responses to narrowband light stimulation were considered for three different durations of light stimuli: transient effects (light onset) and 10s, during the pure visual paradigm, and 50s sustained effects during the visual-cognitive paradigm.

At light onset, activations of the primary visual cortex were detected in both groups for both light conditions, but with a greater extent in HC. Significantly higher response was detected in HC compared to LHON patients under blue light. No significant differences were detected when blue and red light were compared in either groups, and no significant difference in blue vs. red light were detected across groups (Figures 1 and 2, Supplementary Table 1).

**Figure 1.**
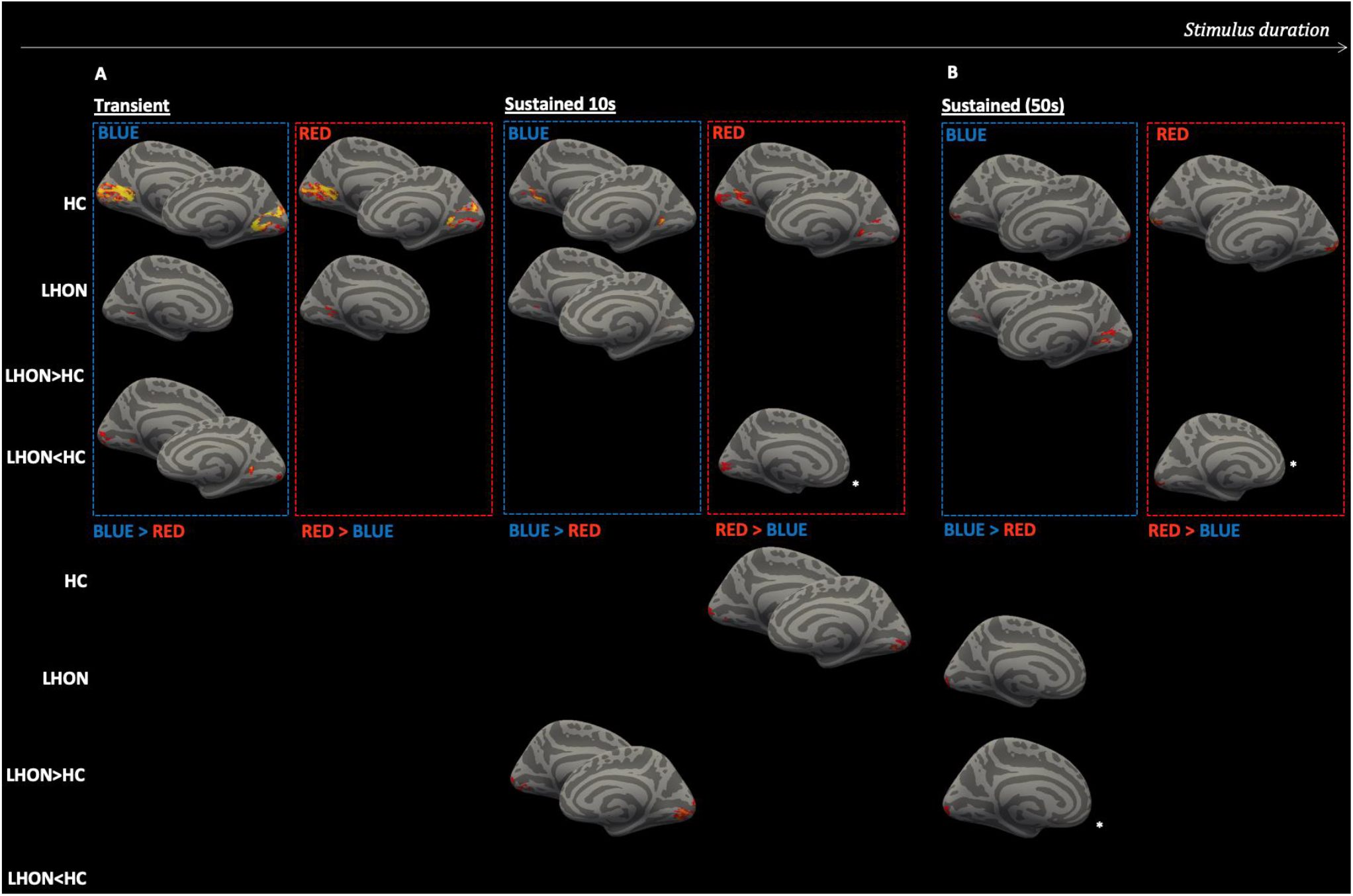
Brain responses to light stimulations. Significant (TFCE-corrected p<0.05) results for LHON and HC brain response to transient effects at light onset (A, left), 10s sustained effects (A right) and 50s sustained effects (B). A: light stimulation effects from the pure visual paradigm; B: light stimulation effects from the visual-cognitive paradigm (the contribution of the cognitive task being regressed out). For the contrasts that gave no significant results, no brain images are shown. Only for the visualization, the results were registered and projected onto freesurfer fsaverage brain surface (left hemisphere on the right). (LHON: Leber’s hereditary optic neuropathy; HC: healthy controls; *: only for visualization purposes clusters are shown at p<0.1 (clusters were however found at p<0.05, see Supplementary Table 1).

**Figure 2.**
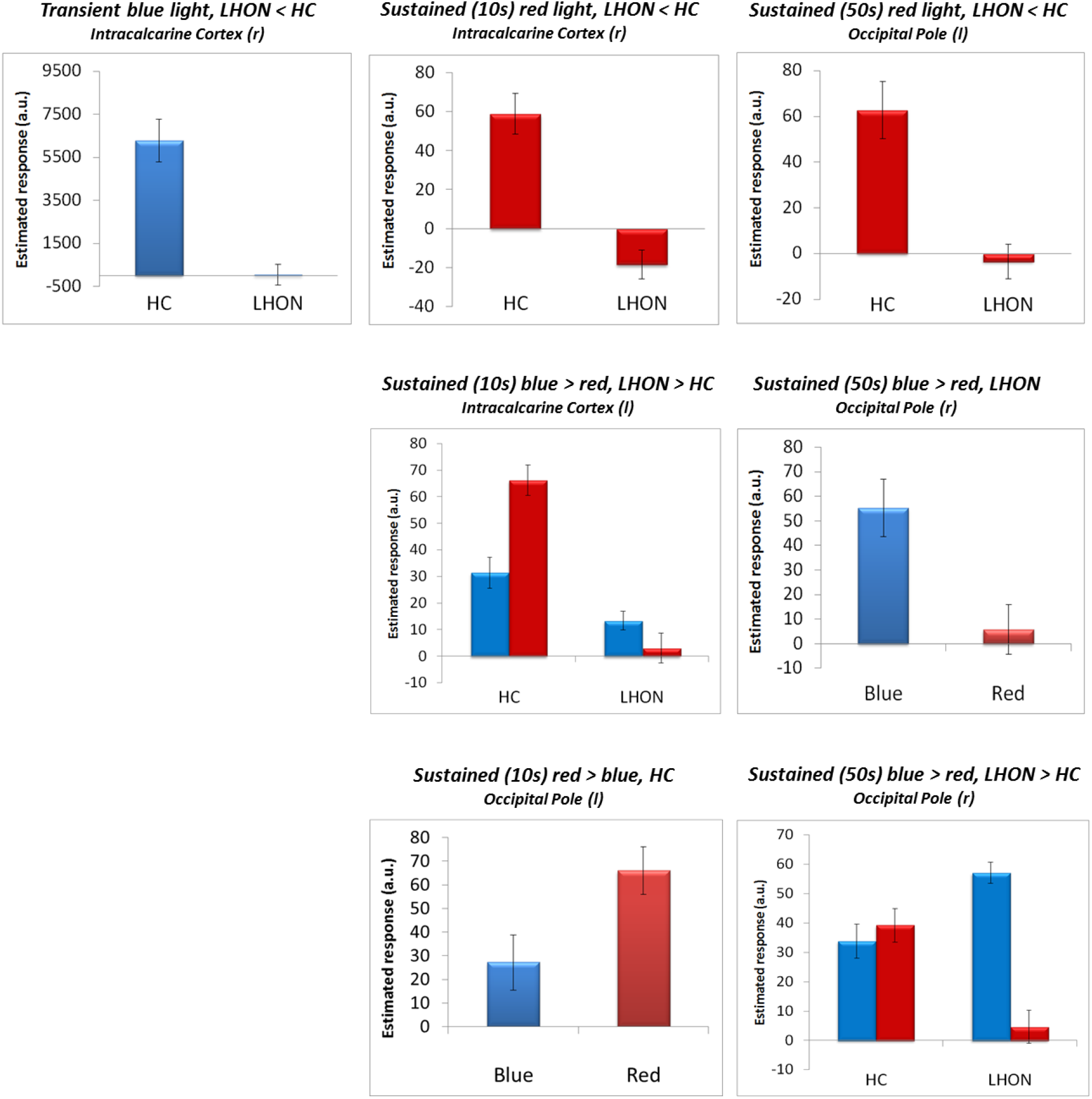
Bar plot of brain responses to light stimulations. Bar plots describing the mean parameters estimates of the significant voxels that were found for the comparisons between light conditions and/or groups (average in arbitrary units ± standard error of the mean). A representative brain response, taken from the main significant cluster, is displayed for each contrast yielding a significant difference.

Both groups showed sustained responses to 10s and 50s exposure to blue light over several parts of the primary visual cortex. In contrast, sustained visual cortex responses to 10s and 50s red light exposure were only detected in HC. Accordingly, sustained responses to both 10 and 50s red light were significantly higher in HC than in LHON patients.

When assessing the interaction between light conditions and groups, sustained responses were greater under blue vs. red light exposure in patients relative to HC in the occipital cortex for both 10 and 50s (Figures 1 and 2, Supplementary Table 1).

No significant correlations were found in LHON patients between functional visual responses under either blue or red light and ophthalmological data, namely visual acuity, visual fields and retinal nerve fibre layer thickness.

### Light modulation of cognitive brain responses

Executive brain responses, isolated by subtracting 0-back brain responses from 3-back responses, were observed in the typical brain areas sustaining working memory (Collette et al., 2006) and similar between the two groups, and encompassed the prefrontal, parietal and cingulate cortices, thalamus, and putamen (Supplementary Figure 1; Supplementary Table 2). No group differences were detected suggesting that both patients and HC successfully and similarly performed both tasks. We then examined whether executive brain responses were affected in a wavelength-dependent manner. Analyses reveal that, compared to red light exposure, blue light exposures increased executive brain responses in LHON patients in the middle frontal gyrus (Figure 3, Supplementary Table 3). No such a significant difference was detected in HC and groups did not significantly differ when considering the differential impact of light wavelength on executive responses.

**Figure 3.**
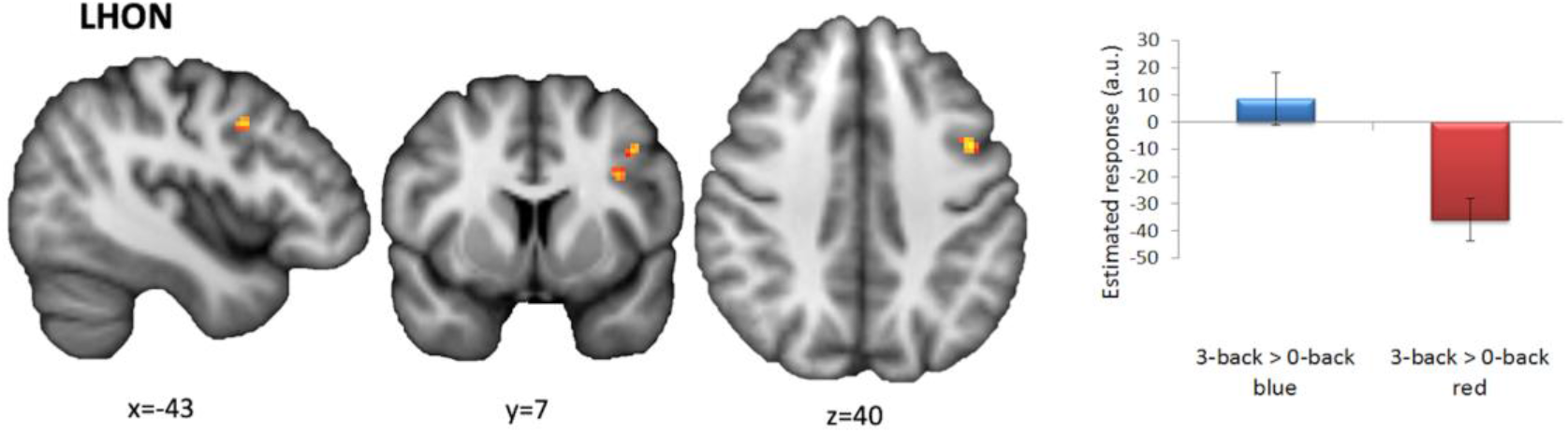
Brain response in LHON patients is modulated by light conditions during the attentive task. Significant (TFCE-corrected p<0.05) results for brain responses to the interaction between 3-back task and different light conditions. Results are shown only for LHON group effect since other contrasts in HC gave no significant results. The background image is an average of individual T1-w scans - in radiological convention. Only for visualization purposes the cluster is shown at p<0.1 (it was however found significant at p<0.05, see Supplementary Table 3). The bar plot describing the mean parameters estimates is reported in the lateral panel.

In addition, no significant correlations were found in LHON patients between functional brain cognitive responses under either blue or red light and ophthalmological data, namely visual acuity, visual fields and retinal nerve fibre layer thickness.

## DISCUSSION

This study investigated the light-driven modulation of brain activity by mRGCs through fMRI in a cohort of chronically and severely affected LHON patients, large enough to allow for group level statistical inferences (and therefore effect generalization). LHON provides an unparalleled opportunity to test the visual impact of mRGCs on the human brain, as the neurodegenerative process substantially ablates the general population of RGCs that contribute to formed vision - without affecting rods and cones – while leaving mRGCs relatively preserved.

In line with our hypotheses, the results first demonstrated significantly higher primary visual cortex sustained activation in response to blue compared to red light stimulation in LHON compared to HC. In particular, we found V1 cortex activation with both blue and red light at all stimuli durations (transient, sustained 10s and 50s) in HC subjects, whereas for LHON V1 activation was evident only in response to blue light. It appears therefore that our findings arise from a greater relative difference between light conditions (blue vs. red) in LHON patients compared with HC. In addition to recordings of brain activity related to light exposure, our protocol also investigated whether mRGC photoreception would affect an ongoing cognitive brain activity by including an auditory working memory task in one of the fMRI sessions. Interestingly, and as anticipated, executive brain responses were differentially affected by light wavelength, with blue light associated with higher activations than red light, in LHON patients over the lateral prefrontal cortex (or middle frontal areas) typically involved in higher executive function (Otero and Barker, 2014).

A few studies investigated the NIF impact of light without simultaneous cognitive task completion (Hung et al., 2017; McGlashan et al., 2018) and they did not isolate activity over the occipital cortex. The present findings strongly suggest that the output of the remaining mRGCs, most sensitive to blue wavelength, is also including the occipital cortex implying a possible role in visual functions. These findings also support previous studies pointing to a relative sparing of mRGCs in LHON (La Morgia et al., 2010; Moura et al., 2013) and support the inference that the mRGC signal indirectly feeds to the brain cortex mostly devoted to vision in humans, in addition to their classical role in circadian photoentrainment and other NIF functions (Hankins et al., 2008; Vandewalle et al., 2013, 2018; Spitschan et al., 2017). Overall our data imply that mRGC signal modulates occipital activity, potentially contributing to visual function in human beings. Since our LHON patient sample was normal and comparable to the HC sample, except for their visual dysfunction, these findings are indeed going beyond the particular cases of LHON patients.

Other evidences supporting the role of mRGCs in cortical visual processes (Sonoda and Schmidt, 2016) include a direct retinofugal projection of mRGCs to the LGN that, in turn, projects to the primary visual cortex (V1) in mice (Hattar et al., 2006; Hatori et al., 2008; Ecker et al., 2010), rats (Langel et al., 2015) and non-human primates (Dacey et al., 2005; Hannibal et al., 2014).

Furthermore, neurophysiological studies in mice suggest that mRGCs can support spatial visual perception (discrimination of very coarse patterns) in animals lacking the classical rod-cone outer retinal system (Ecker et al., 2010). These physiological studies point to a sustained and scalable response to light stimulation mediated by the dorsal LGN (dLGN) (Brown et al., 2010) in photopic conditions (Davis et al., 2015; Mouland et al., 2017). Melanopsin RGCs may drive a generalized increase of dLGN excitability, conveying information about changing background light intensity and increasing the signal/noise for fast visual responses (Storchi et al., 2015). mRGC projections to the LGN may help in the encoding of visual images by increasing the thalamic representation of scenes in reference to total radiance (Allen et al., 2017). Moreover, knockout mice for melanopsin show an impoverished coding of natural scenes suggesting the influence of mRGCs on the spatial and temporal tuning of dLGN neurons (Allen et al., 2014).

Melanopsin RGCs also contribute to visual processing through the maintenance of the pupil light reflex and light avoidance behaviour (Johnson et al., 2010). Finally, a melanopsin system contribution to brightness discrimination has been demonstrated in mice with and without retinal degenerations (Brown et al., 2012). Psychophysical experiments in healthy human subjects have shown a similar role in brightness perception (Brown et al., 2012; Zele et al., 2018a) and suggested the mRGC capacity to signal slowly changing stimuli of light colour (Zele et al., 2018b). Further support for the contribution of melanopsin to human vision is provided by recent evidence that spatial patterns that were spectrally indistinguishable for cones but had contrast for melanopsin could be discriminated by healthy human subjects (Allen et al., 2019). Likewise an fMRI study in four healthy subjects demonstrated that high contrast melanopsin-specific light stimuli elicited a response in the primary visual cortex, associated with a brightening of visual perception (Spitschan et al., 2017). The class of mRGCs that more likely play a role in visual forming functions is represented by the M4 subtype, which are most similar to the conventional RGC subtype by being highly sensitive to contrast (Schmidt et al., 2014). Specifically, melanopsin photosensitivity contribution of M4 cells output is particularly important for contrast sensitivity functions (Schroeder et al., 2018). It is therefore possible that the blue light-induced occipital activity we report in LHON arise from M4 subtype.

Melanopsin-mediated modulation of cognitive brain activity was previously found in sighted subjects over the same lateral prefrontal cortex areas we isolate in LHON patients (Vandewalle et al., 2007). However, we did not find significant differences between executive responses under blue and red light periods in HC as well as no significant difference between groups. The fact that we found a significant difference in the comparison between blue and red light only in LHON patients but not in HC is presumably due to the higher ratio mRGCs/RGCs reported in LHON. The absence of group differences and light condition difference in HC arises in our view from 2 main factors: i) the smaller sample size, although within good practice suggestion for fMRI studies (Desmond and Glover, 2002) (previous studies in sighted individual included up to 16 volunteers (Vandewalle et al., 2011; Chellappa et al., 2014)), and ii) the reduced magnetic strength (1.5T vs. 3T), leading to a lower signal and SNR and time required for a brain volume acquisition (3s vs. ~2s) (Vandewalle et al., 2007, 2011, 2013). We further emphasize that, despite all these limitations, we were able to isolate a light condition impact while performing a cognitive task in part of our sample (and across both groups without cognitive tasks – cf. above). The fact that differential impact of light wavelength on ongoing brain activity was most evident in the LHON group gives further support to a predominant role of mRGCs in modulating ongoing cognitive activity. Aside from a maximal sensitivity to blue light compared to other wavelengths (Vandewalle et al., 2007, 2013) a similar result was previously suggested in a study in sighted healthy young volunteers in which prior light history was manipulated to affect mRGC sensitivity (Chellappa et al., 2014), and in 3 totally blind subjects due to outer retinal disorders, with no conscious vision but retained NIF photoreception (Vandewalle et al., 2013).

As it is challenging to isolate mRGCs in normal vision (Allen et al., 2019), we cannot exclude a contribution from residual RGCs to our findings and therefore from rods and cones, which are preserved in LHON. However, we observe differences between light conditions and/or LHON and controls in terms of brain activation for longer duration stimuli, which is compatible with a melanopic signature, as opposed to the typically transient response of classical photoreceptors.

Finally, despite brain response modulations by blue light in LHON, we did not find an effect of light on behavioural performances. This is not unexpected, given that, as in previous studies in healthy subjects (Vandewalle et al., 2007), we were careful at keeping task blocks short to avoid any behavioural effects that could contaminate the results. Both patients and controls are cognitively intact and our light stimulation scheme included short exposures to light (< 1 min), which differs markedly from what is described for other investigation meant to trigger improvements in cognitive performance, i.e. hour long exposures, sometimes repeated over a week (Mitolo et al. 2018).

To our knowledge, the present study is the first to explore the effect of light on human brain activity in subjects affected by an inherited optic neuropathy, in particular in LHON that is characterized by a selective relative sparing of the mRGC system and support the idea that mRGC specifically activate the occipital cortex in LHON patients, even when the brain is not engaged in a cognitive challenge, and the prefrontal cortex when engaged in a cognitive process. In conclusion, these results support an indirect role of mRGCs for both NIF and visual forming functions in humans and in LHON patients in particular, opening potential windows for therapy in these patients.

## MATERIALS AND METHODS

### Subjects

Twelve patients with LHON and twelve age-matched controls participated to the study. Patients were consecutively recruited at the Neuro-ophthalmology Clinic, IRCCS Istituto di Scienze Neurologiche di Bologna, UOC Clinica Neurologica, Ospedale Bellaria, Italy. Healthy control (HC) subjects were recruited on a volunteer basis among Hospital and University co-workers. Local Ethical Committee approved the study (EC reference ID #14004), according to the Declaration of Helsinki, and all the participants gave their written informed consent.

Inclusion criterion for patients was a genetically confirmed diagnosis of LHON. Exclusion criteria for both patients and HC were contraindications to MR examination, neurological or psychiatric diseases, use of drugs acting on central nervous system or on sympathetic and parasympathetic system and excessive caffeine (> 4 cups/day) or alcohol (> 14 units/week) consumption; we also excluded volunteers who were shift-workers during the previous year, or had travelled through more than one time zone during the previous 2 months. Other exclusion criteria for HC were ocular hypertension, lens opacity, retinal or optic nerve diseases including macular degeneration and colour vision abnormalities. The Morningness-eveningness questionnaire was used to assess subjects’ chronotype (Horne and Ostberg 1976). The Pittsburgh Sleep Quality Index Questionnaire (pathological score >5) (Buysse et al., 1989) and the Berlin questionnaire (Netzer et al., 1999) were used to assess the presence of sleep disturbances, and the Epworth Sleepiness Scale for excessive daytime sleepiness (ESS≥11) (Vignatelli et al., 2003). Beck Anxiety Inventory (Beck et al., 1988) and 21-item Beck Depression Inventory scales (Beck et al., 1961) were used to evaluate anxiety and depression levels in the study cohort (pathological score ≥14).

### Study Design

#### Before fMRI sessions

During the week preceding the fMRI session, participants were asked to follow a regular sleep schedule (maintaining their habitual sleep routine, with a tolerance interval of 1 hour), to be reported in sleep diaries, and they were also asked to refrain from caffeine, alcohol or other substances acting on central nervous system for 3 days before the MR session. Moreover, participants were trained to the cognitive task administered inside the MR scanner (see below; Training 1, Figure 4-A).

**Figure 4.**
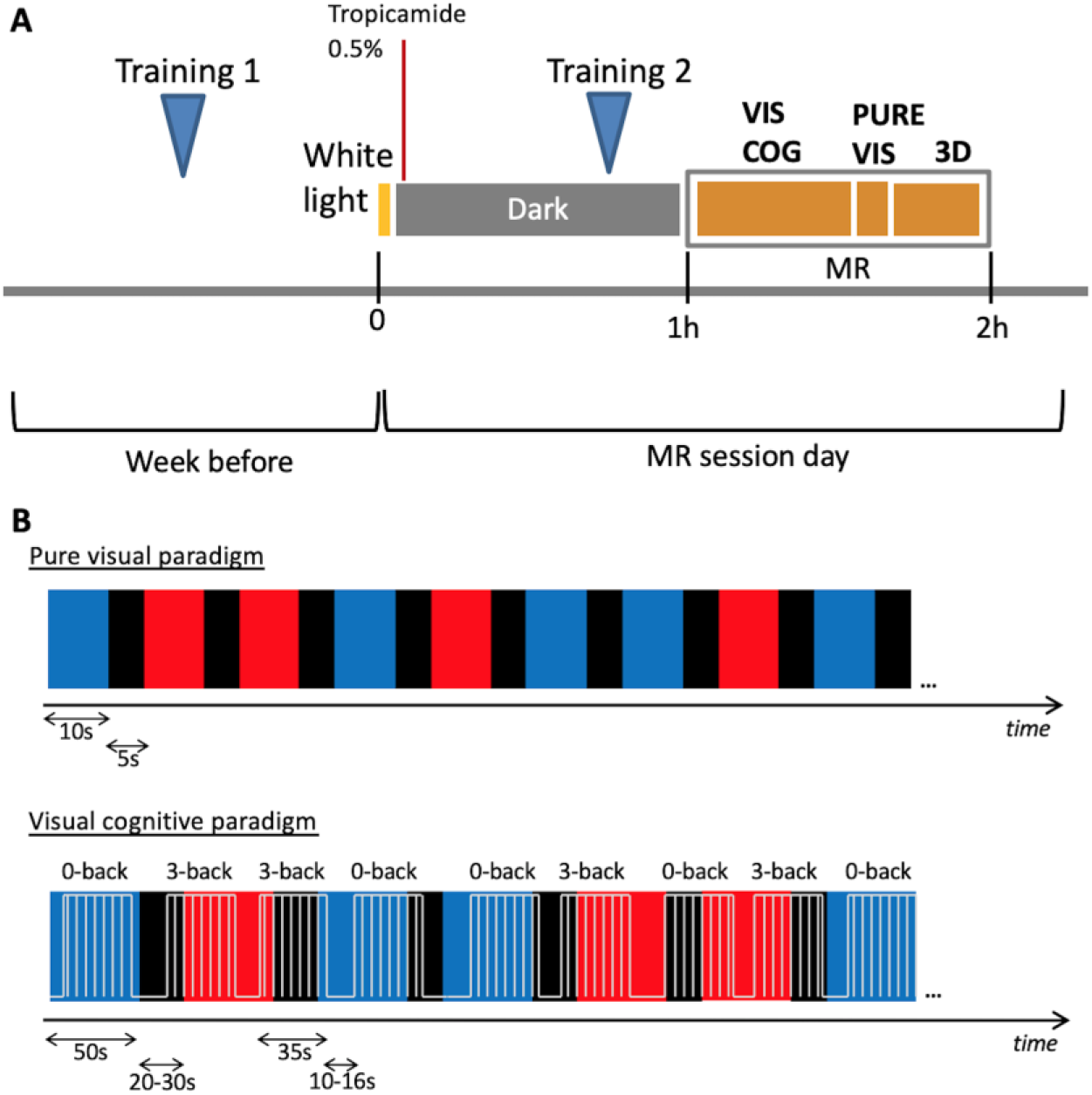
Experimental protocol (A) and schematic representation of fMRI paradigms (B). An example of a sequence of light stimulation (red or blue) is provided, together with the cognitive tasks in the lower display. For representation purposes, time axes are not in scale. (VIS COG: visual cognitive paradigm, PURE VIS: pure visual paradigm, 3D: volumetric structural image).

#### fMRI session

For all participants, acquisitions were performed 4 hours after habitual wake time. Since the seasonal variation in environmental light at the time of acquisition may affect cognitive brain activity (Meyer et al., 2016), the average number of hours of light per-day at the time of MR session for each subject (data from Bologna Guglielmo Marconi Airport weather station, monthly average) was taken into account in all analyses.

On the experimental day, subjects were first exposed to white light (1000-1500 lux) for 5 minutes upon arrival, in order to standardize photic history across participants and level out this potential bias (Chellappa et al., 2014), and 1 or 2 drops of tropicamide 0.5% were administered to both eyes to induce mydriasis and cycloplegia. The subjects were then blindfolded and stayed in a dark room for one hour before the fMRI acquisitions. During the dark adaption, subjects underwent a short second training to the cognitive task (Training 2, Figure 4-A).

#### Light exposure

Narrow interference band-pass filters were used to produce both narrowband illuminations: blue - 480nm (Full width at half maximum (FWHM): 10nm)- and red - 620nm (FWHM: 10 nm). The blue wavelength was meant to correspond to melanopsin maximal sensitivity, while the red light was equally away from the peak sensitivity of the photopic visual system (i.e. 550nm), while being close to undetected by mRGCs.

A filter wheel (AB301-T, Spectral Products, NM) was computer-controlled to switch band-pass filters and thereby change light wavelength. The light was transmitted by a metal-free purpose-built optic fibre (Fiberoptics Technology Inc, CT) from a source (DC951H illuminator, EKE lamp, Dolan-Jenner) to two small diffusers placed in front of the subjects’ eyes (Ground glass diffuser 220 Grit, Thorlabs). Diffusers were designed for the purpose of this study and ensured a reasonably uniform illumination over the visual field; they were placed approximately 2 cm away from subjects’ eyes. Irradiance could not be measured directly in the magnet, but the light source was calibrated and photon flux estimated to be 5×10^13^ph cm^−2^s^−1^ (Power meter PM100D, Thorlabs with Silicon Power head S120VC), corresponding to an irradiance of 20.7 μW/cm^2^ for the blue light and 16.0 μW/cm^2^ for the 620nm red light On the lux scale, to quantify the effective illuminance for human photopigments (Lucas et al., 2014), for a wavelength of 480nm and an irradiance of 20.7 μW/cm^2^ we obtained the following values: photopic illuminance = 19.87 lux, melanopic illuminance (mRGCs) = 165.01 lux, rhodopic illuminance (rods) = 118.28 lux, cyanopic illuminance (S-cones) = 78.87 lux; chloropic illuminance (M-cones) = 63.1 lux, erythropic illuminance (L-cones) = 32.97 lux.

For a wavelength of 620nm and an irradiance of 16.0 μW/cm^2^ we instead obtained: photopic illuminance = 41.71 lux, melanopic illuminance (mRGCs) = 0.13 lux, rhodopic illuminance (rods) = 0.97 lux, cyanopic illuminance (S-cones) = 0 lux; chloropic illuminance (M-cones) = 16.37 lux, erythropic illuminance (L-cones) = 51.92 lux.

The light device produced no perceptible sounds or temperature change. The total amount of blue light received during the experiment was 4 orders of magnitude below the blue-light hazard threshold as defined by the International Commission on Non-Ionizing Radiation Protection (ICNIRP Guidelines 2013).

#### fMRI paradigms

The first paradigm tested was meant to investigate the possible role of mRGCs in a pure visual setting. Participants were exposed to blue or red lights for periods of 10s separated with 5s of complete darkness (<0.01 lux), with a random colour alternation, for a total duration of 5 minutes (Figure 4-B).

In order to investigate mRGC-driven modulation of brain responses during a working memory task, a cognitive paradigm was constructed based on previous studies (Vandewalle et al., 2007, 2013) (Figure 4-B). The paradigm included 50s illumination periods under blue or red light exposure, separated by dark periods of 20 to 30s (mean 25s). While exposed to light or maintained in darkness, participants performed 35s blocks of either 0-back and 3-back auditory task separated by rest periods lasting 10 to 16s (mean 13s). Both auditory tasks consisted in series of consonants. The 0-back task was a simple letter detection task during which subjects were requested to state whether or not the consonant was an “r”. The 3-back task is a working memory task requesting to state whether each consonant was identical to the consonant presented three stimuli earlier. It is an executive task probing maintenance and updating of information as well as attention and auditory processing (Cohen et al., 1997, Collette et al., 2006).

Responses were given by pressing a button on a MR-compatible handgrip when the answer was yes. Stimuli consisted of nine Italian monosyllabic consonants (duration = 0.5 s, Inter-Stimulus Interval= 2 s), produced using COGENT 2000 (www.vislab.ucl.ac.uk/cogent.php), implemented in MATLAB (MathWorks, MA), and transmitted to the participants using MR compatible headphones. Series of stimuli were constructed with 30% hits so that the difficulty level was similar in all blocks, were presented only once and were randomly assigned to a task block. Each auditory task block consisted of a series of 14 consonants. A total of 42 blocks were presented, 21 of 0-back and 21 of 3-back, randomly alternated. Each type of task was preceded by a short vocal instruction. The cognitive task was totally uncorrelated to the light condition, i.e. presentation of task blocks was independent from light changes, so that both the impact of light on prefrontal cognitive brain activity and occipital visual brain activity could be investigated separately. The duration of the cognitive paradigm was about 35 minutes.

### fMRI acquisition

fMRI acquisitions were performed with a 1.5 T system (GE Medical System Signa HDx 15), equipped with an 8-channel brain phased array coil. The static magnetic field of the apparatus was therefore lower than previous 3T fMRI studies on the NIF impact of light (Vandewalle et al., 2007, 2011, 2013). Since signal and signal-to-noise ratio (SNR) decrease non-linearly as a function of magnetic field, this implies that sensitivity of the apparatus was much lower than previously. Yet, the excellent access to the rare phenotype of interest at the University of Bologna, i.e. relative increase in mRGC photoreceptionin LHON patient, led us to postulate that the most prominent effects, i.e. the greater relative difference in mRGC/RGC output, would be detectable with the 1.5T apparatus.

Functional MR images were acquired with a multislice T2*-weighted gradient-echo-planar sequence using pure axial slice orientation (34 slices, thickness 4 mm, in-plane resolution 1.875×1.875 mm, field of view FOV=240×240 mm, matrix size=98×98×34, repetition time TR=3000 ms, echo time TE=40 ms, flip angle=90°). High-resolution volumetric structural images were acquired using a T1-weighted fast spoiled gradient echo (FSPGR) sequence, (TR=12.4 ms, TE=5.2 ms, inversion time TI=600 ms, flip angle=10°, matrix size=256×256 mm, FOV=256×256 mm, voxel size 1×1×1 mm). Acquisitions started with the visual cognitive paradigm, then the pure visual paradigm followed, and the structural images acquisitions.

### fMRI data analysis

Analyses of fMRI data were performed with the FSL software (https://fsl.fmrib.ox.ac.uk/fsl/). Image pre-processing included motion correction through rigid body registration (MCFLIRT, Motion Correction FMRIB’s Linear Image Registration Tool), high-pass filtering (cut-off 100s for pure visual paradigm and 150s for visual-cognitive paradigm), spatial smoothing (gaussian kernel FWHM 5mm) and slice timing correction.

At the single subject level, changes in brain responses were estimated by using a general linear model, in which aspects of interest were modelled using boxcar or stick functions convolved with a double-gamma hemodynamic response function. In particular, for the pure visual paradigm, the following explanatory variables (EV) were included in the design matrix: blue and red (modelled with boxcar functions), blue on, blue off, red on and red off (modelled with stick functions).

Movement parameters derived from realignment for motion correction were added as covariate of no interest. COPE (Contrast of Parameter Estimates) maps were generated for the following contrasts: blue, red, blue > red, blue < red, blue on, red on, blue on > red on, blue on < red on. Light offsets were included as regressors of no interest while transient effects in brain activity induced by light onset were considered as effect of interest.

Regarding the visual cognitive paradigm, boxcar functions were used to model 0-back task blocks, 3-back task blocks, blue illumination periods and red illumination periods. Stick functions were used for light onset and offset which were considered as covariate of no interest together with movement parameters. The following EVs were included in the design matrix: 0-back, 3-back, blue, red (modelled with boxcar functions), blue on, blue off, red on, red off (modelled with stick functions), and then the interactions between light and task: 0-back under blue, 0-back under red, 3-back under blue, 3-back under red. In all contrasts, executive brain responses were isolated by subtracting brain responses to the 0-back task from the brain responses to the 3back task. We assessed these brain responses irrespective of the light condition and then evaluated the impact of light on executive responses. COPE maps were generated for the following contrasts: 3-back > 0-back, blue, red, blue > red, blue < red, (3-back blue – 0-back blue) > (3-back red – 0-back red), [(3-back blue – 0-back blue) < (3-back red – 0-back red)].

Functional images were linearly aligned to structural images and structural images were non-linearly aligned to the MNI template. At the group level, comparisons between LHON patients and healthy controls (HC) were carried out with non-parametric statistics obtained by permutation methods (FSL randomise, with 5000 permutations). Age, sex and the average numbers of hours of light per day at the moment of MRI acquisitions were added as covariate of no interest. Comparisons were performed within pre-defined regions of interests: primary visual cortex for the visual effects and prefrontal brain regions associated with working memory tasks for the visual-cognitive effect. Precisely, V1 was defined based on Juelich histological atlas (Eickhoff et al., 2005) definition at 25% probability, while regions involved in working memory task were defined according to a recent meta-analysis results (Wang et al., 2019) by drawing a sphere of 10mm radius around each coordinate reported for all active main effect and load condition. Statistical inferences were made from statistical maps that were corrected for multiple comparisons with a threshold free cluster enhancement (TFCE) method, considering significant results at p<0.05. An analogous approach was used to investigate possible correlations between fMRI results and patients ophthalmological data.

### Demographic and behavioural data analysis

Normal distribution of all data types was checked with a Shapiro-Wilk test. Gender was compared between the two groups with Pearson’s χ^2^ test, while age and the average hours of light were compared with a t-test. The performances in the two training sessions of the n-back cognitive tasks were compared between sessions with a paired t-test, and between patients and controls with a t-test. As for the performance of the cognitive task during MR acquisitions, a two-way mixed design ANOVA was performed, with group (patients or controls) as independent factor and light conditions (blue, red, darkness) as the three-level repeated measures.

### Ophthalmological evaluations

Both patients and controls performed an ophthalmological evaluation which included the assessment of visual acuity (ETDRS chart), measurement of intraocular pressure (IOP), evaluation of the anterior chamber by means of slit lamp and of the fundus oculi by means of direct ophthalmoscopy. Moreover, participants performed evaluation of colour vision (Ishihara’s Test for Colour-Blindness-Kanehara Shupman Co., Tokyo, Japan), computerized visual field (Humphrey, Zeiss) and optical coherence tomography (OCT) (Stratus, Zeiss). For correlation analysis the following metrics were used: visual acuity, mean deviation for computerized visual fields, retinal nerve fibre layer (RNFL) thickness average and single quadrants (temporal, superior, nasal and inferior) thickness (for more details on OCT methods see Barboni et al., 2010).

## Supplementary material

Supplementary file 1: Supplementary Figure 1. Significant (TFCE-corrected p<0.05) group-level results for responses to 3-back task compared to the control condition (0-back) irrespectively of light conditions.

Supplementary file 2: Supplementary Table 1. Brain response to monochromatic light stimulation.

Supplementary file 3: Supplementary Table 2. Cluster data for group-level results for responses to 3-back task compared to the control condition (0-back) irrespective of light condition.

Supplementary file 4: Supplementary Table 3. Brain response in LHON patients is modulated by light conditions during the attentive task.

## Acknowledgments

We thank Fondazione del Monte di Bologna e Ravenna for the financial support. It had no role in study desigg, in the collection, analysis and interpretation of data, in the writing of the report and in the decision to submit the article for publication.

